# Polymer additives to personal protective equipment can inactivate pathogens

**DOI:** 10.1101/2021.04.01.438151

**Authors:** Alan B. Dogan, Katherine Dabkowski, Horst A. von Recum

## Abstract

Face masks have been proven to be medicine’s best public health tool for preventing transmission of airborne pathogens. However, in situations with continuous exposure, lower quality and “do-it-yourself” face masks cannot provide adequate protection against pathogens, especially when mishandled. In addition, the use of multiple face masks each day places a strain on personal protective equipment (PPE) supply and is not environmentally sustainable. Therefore, there is a significant clinical and commercial need for a reusable, pathogen-inactivating face mask. Herein, we propose adding quaternary poly(dimethylaminohexadecyl methacrylate), q(PDMAHDM), abbreviated to q(PDM), to existing fabric networks to generate “contact-killing” face masks – effectively turning cotton, polypropylene, and polyester into pathogen resistant materials. It was found that q(PDM)-integrated face masks were able to inactivate both Gram-positive and Gram-negative bacteria in liquid culture and aerosolized droplets. Furthermore, q(PDM) was electrospun into homogeneous polymer fibers, which makes the polymer practical for low-cost, scaled-up production.

**Graphical Abstract:** 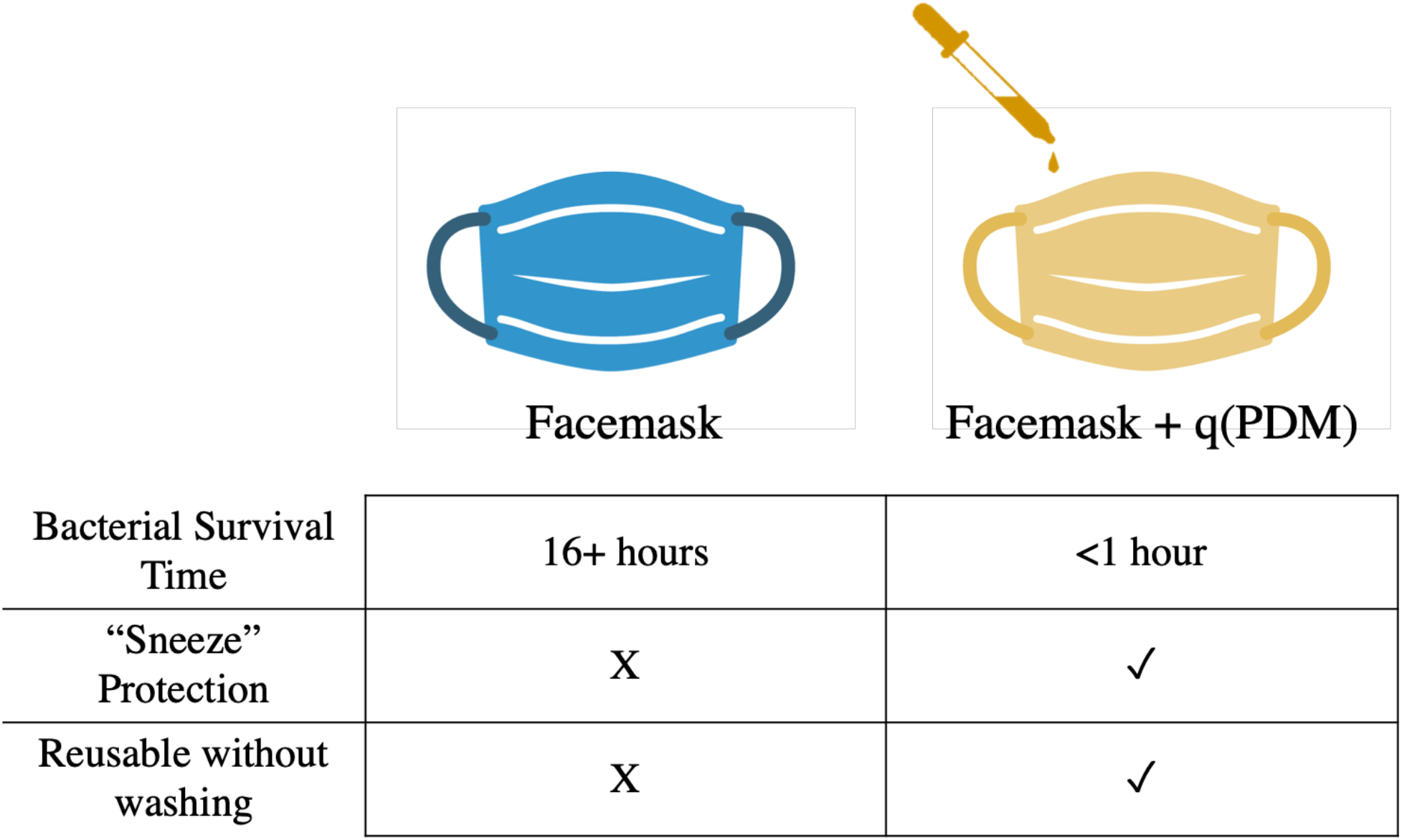

## 1. Introduction

The COVID-19 pandemic has shown that despite significant advances in healthcare and biotechnology, face masks are still the most impactful public health tool available for preventing transmission of airborne pathogens. Surgical masks have shifted from being a ‘healthcare product’ to an essential ‘consumer product’, which triggered personal protective equipment (PPE) shortages across the world^1^. While, historically, facemasks take only a few cents to manufacture, due to limited supply and high demand between primary healthcare providers, patients, and everyday people, the price of single-use PPE has risen exponentially, up to $30 per mask in March 2020.

Even the gold-standard of mask PPE, such as N95 respirators and their international counterparts (e.g. KN95), are only effective for preventing transmission with proper sterile techniques and ‘fit tests’ and can only be sterilized up to three times before filtering efficiency decreases^2,3^. Many consumer mask users also lower the efficacy of facemasks by either wearing a mask incorrectly, wearing an improper facemask, or inappropriately reusing facemasks without proper decontamination protocols (based on CDC suggestions)^4,5^. 3D printing of PPE has also become increasingly popular, however the masks produced in this way are not FDA approved and have issues regarding heterogeneity and inefficient filtration capacity^6,7^. Newer antiseptic facemask materials have claims of integrating nanofibers, nanoparticles, copper and metal particles/fibers, heat-producing components, and surfactants to generate an antimicrobial and antiviral effect; however, these are not yet widely available to consumers due to high costs or issues in reproducability^8–13^.

Traditional surgical face mask materials – including polypropylene, polyethylene, cotton and polyester – allow for bacterial and viral adhesion. While this may be insignificant for single use PPE, for consumers who seek to use these masks daily, noncompliance and mishandling of PPE greatly increases risk of transmission and homemade solutions have been found to only filter 10-60% of respiratory-sized droplets^14,15^. This poses a risk for transmission, especially those who are less-dexterous and forgetful. Therefore, there is a current patient and consumer need for a low-cost, reusable face mask that prevents respiratory transmission by contact killing viruses and bacteria while maintaining breathability.

While the need for a contact-killing face mask is obvious in the era of COVID-19, a growing geriatric population, rapid urbanization, and the increasing prevalence of airborne diseases are expected to make the demand for reusable face masks continue to increase^16^.

Our proposed solution is integrating a polymeric quaternary ammonium compound (polyQAC), specifically quaternary ammonium poly(dimethylaminohexadecyl methacrylate), q(PDM), into existing PPE solutions to contact-kill bacteria and viruses to prevent secondary transmission of diseases from PPE mishandling. Quaternary ammonium compounds (QACs) are a class of cationic polymers typically used in food processing and surface sanitation due to their ability to contact-kill bacteria and viruses while reducing protein adhesion to surfaces^17^. While QACs have been utilized for decades in sanitation applications, polymerized QACS (polyQACs) have recently shown to have potent antibacterial and antiviral properties and limited cytotoxicity, making them attractive for biomedical implants, wound dressings, catheters and dental implants^18–20^. Many recent studies regarding these materials cite enhanced therapeutic indices and a lower likelihood of developing antibacterial resistance in comparison to their monomer counterparts^21^.

Herein, we plan to leverage the antibacterial and antiviral activity of polyQACs to augment standard surgical facemasks to ensure sterility over time, thereby allowing the reuse of PPE in both a clinical and consumer setting. Current combination products for contact-killing masks rely on metal nanoparticles, heat, and electrical currents; however, all of these solutions require significant changes in the way we manufacture PPE^22–24^. On the other hand, q(PDM) can be integrated into existing PPE by either 1) spraying or soaking a q(PDM) solution onto a facemask and allowing it to dry or 2) electrospinning fibers of pure q(PDM), which is a relatively low-cost and high-efficiency technique capable of manufacturing large quantities of fibers in a relatively short period of time.

**Figure 1:**
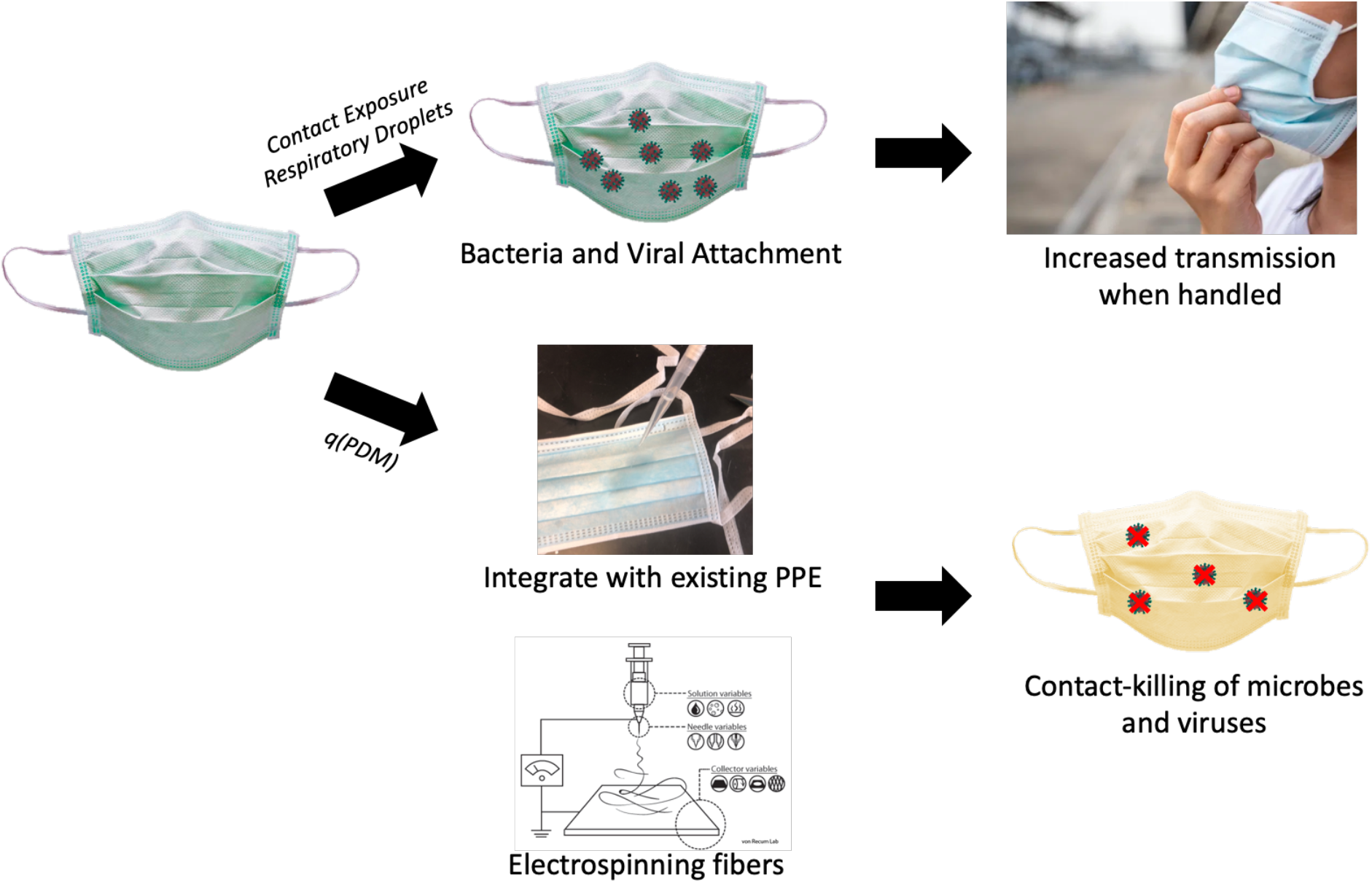
Exposure to airborne pathogens allow for viral and bacterial adhesion to traditional PPE fabrics. Integration of q(PDM) seeks to enable traditional facemasks to exhibit antiseptic properties to ensure PPE remains near-sterile after exposure.

## 2. Materials and Methods

### 2.1 Materials

2-(Dimethylamino)ethyl methacrylate (DMAEMA), 1-Bromohexadecane, and 2,2’-Azobis(2-methylpropionitrile) (AIBN) were purchased from Sigma-Aldrich (St. Louis, MO). Nebulizer parts and precision air compressor were purchased from Shop Nebulizer (Brookfield, CT). All other reagents, solvents, and chemicals were purchased from Fisher Scientific (Hampton, NH) in the highest grade available.

### 2.2 Q(PDM) Synthesis

Q(PDM) was synthesized in a two-step reaction 1) a modified Menschutkin reaction to synthesized a quaternary ammonium methacrylate monomer (q(DMAEMA)) and 2) free radical polymerization of q(DMAEMA) with AIBN **(Figure 2)**. Briefly, 10 mmol of both 1-bromohexadecane and DMAEMA were dissolved in 3 mL of ethanol (EtOH) and reacted at 70°C for 24 hrs under N_2_ atmosphere and stirred. The resulting solution was then distilled under vacuum at 60°C to remove both excess EtOH and residual monomethyl ether hydroquinone (polymerization inhibitor) from the DMAEMA stock. Upon complete removal of solvent an opaque, white solid was observed.

**Figure 2:**
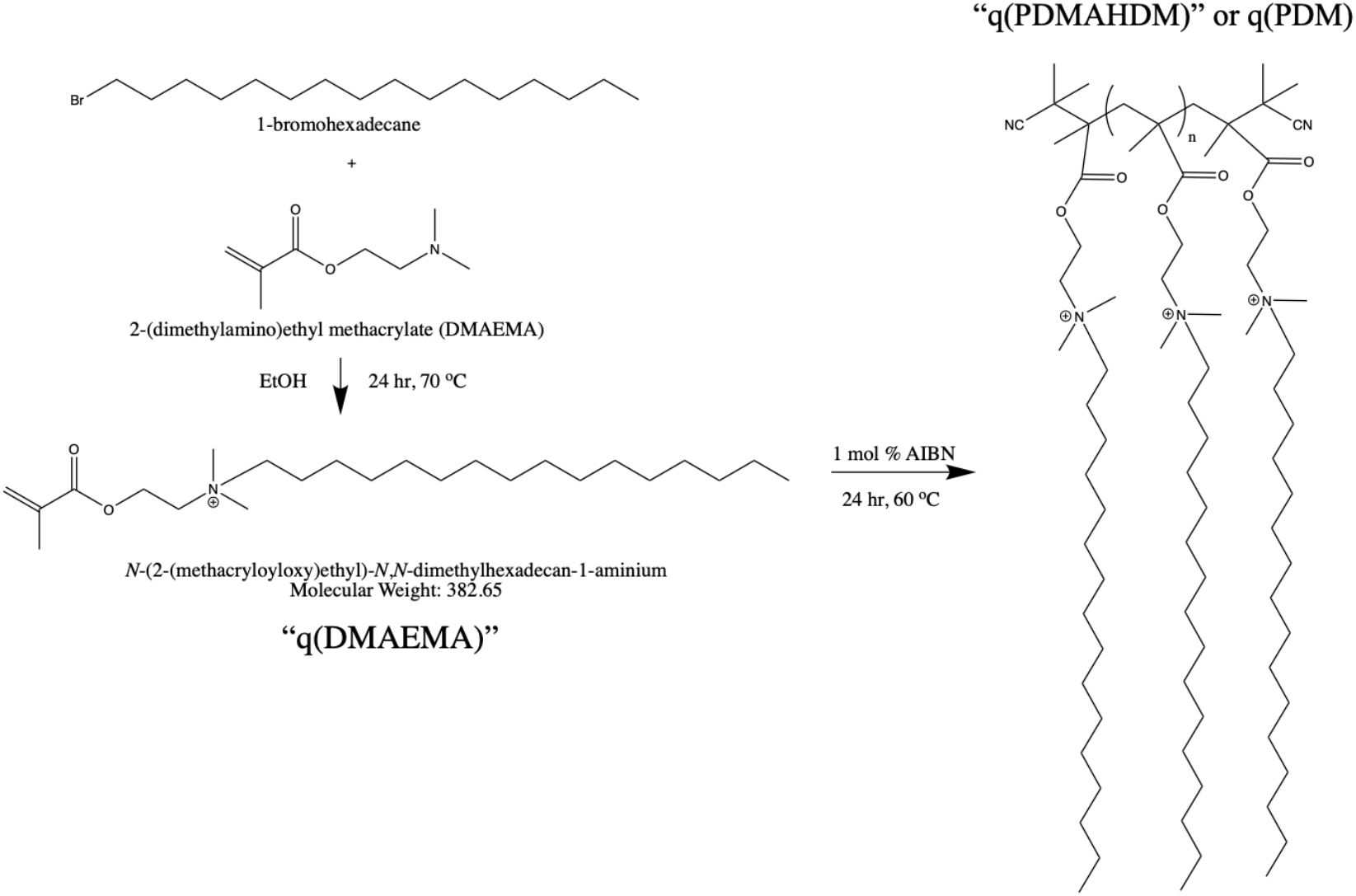
Two-step chemical synthesis of q(PDM) including 1) formation of qDMAEMA monomer via modified Menschutkin reaction and 2) formation of q(PDMAHDM) (abbreviated to q(PDM)) via free radical polymerization

Once distilled, the solution was placed in N_2_ atmosphere, heated to 60°C (stirred), and 1 mole percent AIBN (in ~300μL EtOH) was added to solution and allowed to react for 24 hrs. Upon reaction completion, the mixture was observed to be viscous and a clear/yellow polymer gel was observed. Synthesized polymer was dried and stored at 2°C.

Both one-pot synthesis and quaternization after free radical polymerization were attempted, however sufficient polymerization and functionalization were not achieved in each case, respectively.

### 2.3 NMR and Molecular Weight Estimation

Nuclear magnetic resonance (NMR) was used to verify successful conjugation during synthesis steps. All spectra of presented chemical species were recorded by Bruker 300 MHz NMR system (Bruker, Germany) in DMSO-D_6_. ^1^H-NMR spectra were analyzed on Bruker Topspin software (4.0.9). Unique peak integrals for polymer end groups, backbone, and functionalized alkane groups were used to quantify percent functionalization and molecular weight.

### 2.4 Bacteria-Broth Shake Test

Polymer anti-bacterial properties were assessed with a modified broth-dilution test against both S. aureus and E. coli. 3 mL of ~10^9^ colony forming units (CFU) / mL were incubated for 24 hrs at 37°C with 40-60 mg of dried q(PDM). Before and after incubation, an area scan of each well was taken at 600nm on a Synergy H1 Hybrid Multi-Mode Microplate Reader (BioTek Instruments, Inc., Winooski, VT). Results were reported as normalized percent growth and percent growth reduction per mg polymer, as compared to negative controls.

### 2.5 Q(PDM)-coated Mask Preparation

7.5-20 w/w% solutions of q(PDM) were made in EtOH by mixing dried polymer with solvent on an end-over-end mixer at room temperature, overnight (ON). Polymer solutions were evenly applied to general surgical face masks using pipettes. Solutions were allowed to dry at room temperature and mask mass difference was used to calculate polymer density (mg polymer / mm^2^ mask).

### 2.6 Scanning Election Microscopy (SEM)

SEM sample images were completed at the Swagelok Center for Surface Analysis of Materials at Case Western Reserve University. Samples were mounted on carbon tape, sputter coated with palladium, and imaged on a FEI Helios NanoLab 650. Fiber diameters were determined using ImageJ.

### 2.7 Upright-Cup Permeability Test

Material permeability was assessed by an upright cup test, commonly used for fabric material analysis. ~8-10 mL of water was weighed out into glass scintillation vial and facemask materials were secured with parafilm to form a seal. Samples were incubated at 37°C, 18% humidity, for 24-72 hrs and loss in water mass was recorded.

### 2.8 Nebulized Bacteria Filtration Assay

Modified PPE materials were tested in vitro for filtering and contact-killing efficiency via a modified protocol based on an Andersen cascade impactor assembly. As shown in **Figure 3**, a nebulizer was placed in series with a Precision Medical EasyComp Air Compressor and aerosolized droplets were pushed through the sampled material, with flow-through collected on a sterilized petri dish. Two conditions were tested – 1) ‘chronic exposure’, where bacteria were nebulized without a significant pressure gradient for 10 minutes and 2) ‘acute exposure’, where bacteria was nebulized and exposed to the material surface for 30 seconds with ~2 psi pressure gradient, to simulate a ‘sneeze’ or ‘cough’. Cultured bacteria, either S. aureus or E. coli (~10^9^ CFU/mL), were diluted 1:1 with diH_2_O and loaded into the nebulizer cup and replaced for each trial. The petri dish collecting flow-through (FT) and a swab of the front (F) and the back (B) of the mask was incubated ON at 37°C. Both ‘perfect seals’ and ‘imperfect seals’ (5-10 needle holes poked through the mask before use) were tested.

**Figure 3:**
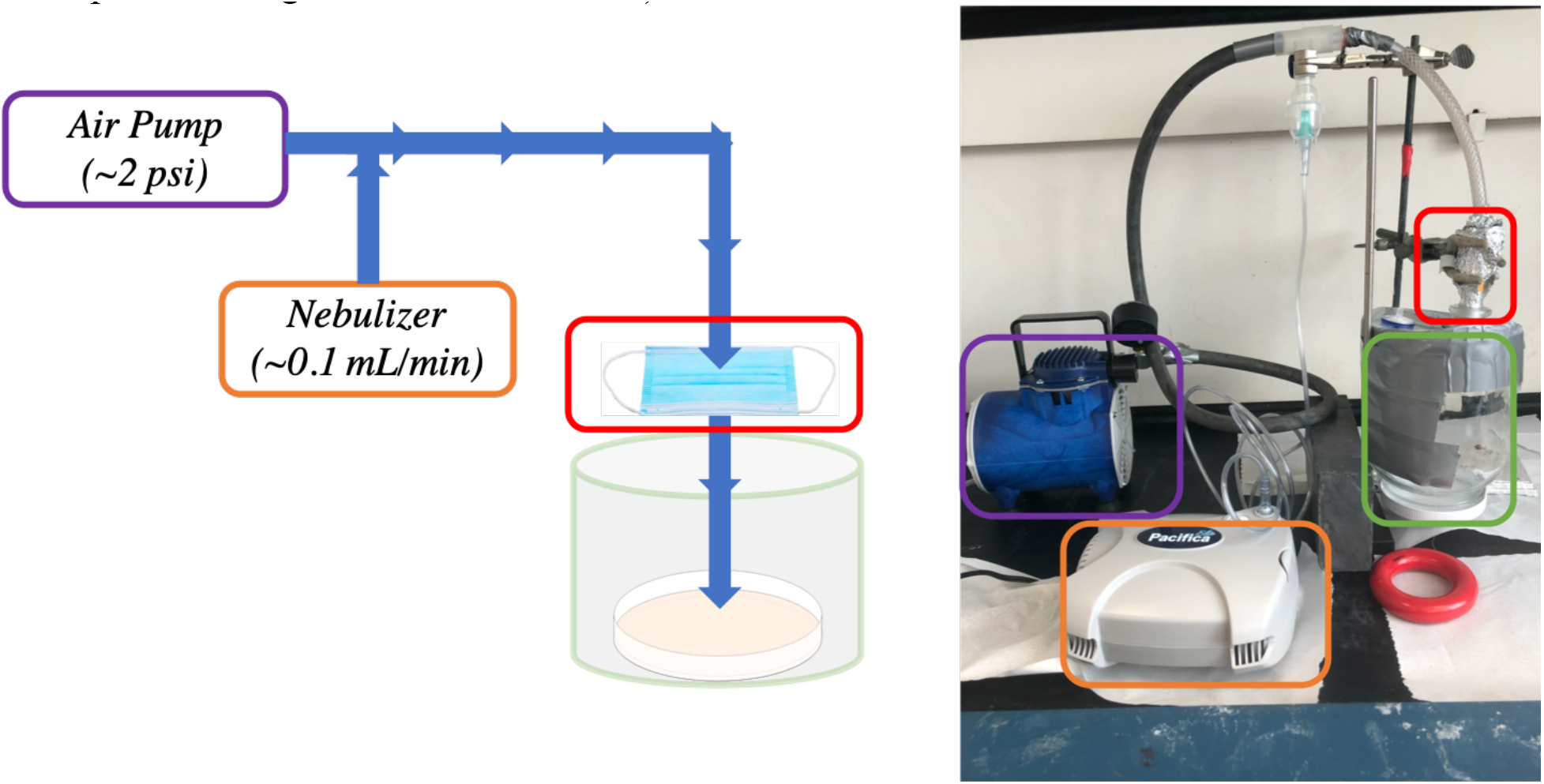
Experimental setup for modified PPE filtering and contact-killing efficiency in vitro. In future work, the petri dish can be replaced by an Andersen cascade impactor for more detailed particle flow-through data.

### 2.9 Surface Contact-Killing Assay for Facemask Fabrics

In order to quantify the modified PPE’s ability to inhibit bacterial growth on its surface, we used a modified surface contact-killing assay. Briefly, 200 μL of diluted confluent bacteria (1:3 dilution in diH_2_O) was applied to the front surface of the facemask and spread. At each recorded timepoint, a sterile scraper was used to sample a small area of the material and applied as a single streak to a petri dish. 24 hours after the sample timepoint, the streak was imaged for CFU analysis.

### 2.10 Electrospinning q(PDM) Fibers

Electrospinning was performed using a Spraybase® (Cambridge, MA) Electrospinning Starter Kit, a 20kV power supply and a syringe pump. Briefly, polymer solutions were loaded into a 3mL syringe and were spun with varied rates (mL/hr), voltage potentials (1-20kV), and solvents (THF, DMF, EtOH). Emitted samples were collected on aluminum foil sheets.

## 3. Results

### 3.1 Q(PDM) two-step synthesis

Q(PDM) synthesis was confirmed via ^1^H-NMR. Confirmation of both free radical polymerization and the successful conjugation of 1-bromohexadecane to DMAEMA backbone was observed by the coexistence of DMAEMA’s hydrocarbon (−CH_2_-) peaks at δ =4.52 ppm, hexadecane’s terminal methyl group (−CH_3_) at δ =1.37 ppm, and AIBN’s two terminal methyl groups (−CH_3_) on the ends of each terminated polymer chain at δ =0.88 ppm. Percent quaternization was determined by comparing peak integrals of hexadecane terminal groups (δ =1.37 ppm; 3 hydrogens) and DMAEMA’s unique ‘-CH_2_-‘ group (δ = 4.52 ppm; 2 hydrogens), yielding approximately 70% quaternization (n=3 batches). Approximate molecular weight was obtained by comparing peak integrals of AIBN’s terminal methyl groups (6 hydrogens per end-group) with DMAEMA’s ‘-CH_2_-‘ group to obtain N number of DMAEMA monomers and N’ number of quaternized DMAEMA monomers. Molecular weight was estimated using the following equation:

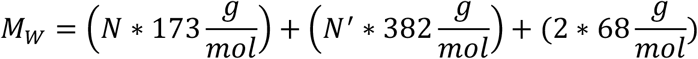

Q(PDM)’s molecular weight ranged between 9,800-30,000 g/mol.

Other polymerization attempts suffered from either insufficient quaternization (polymerize DMAEMA, then quaternized; ~13 kg/mol, ~16% quaternization) or insufficient polymerization (‘one-pot’ synthesis; ~8 kg/mol, 24% quaternization).

### 3.2 Solid q(PDM) polymer contact kills both Gram-Negative and Gram-Positive Bacteria

To test the contact-killing bulk properties of q(PDM), a simple, bacteria-broth shake test was used, wherein the population of viable bacteria greatly outnumbered the total surface area of the polymer. We found that q(PDM) effectively reduced the growth of both S. aureus and E. coli (**Figure 5a**); however, the polymer was found to be more effective against S. aureus than E. coli, which matched previous studies with similar polymers^25^. Q(PDM)’s antimicrobial activity also appeared to be surface-area dependent, as seen in **Figure 5b**, where bacteria were only observed growing in regions without direct polymer contact.

### 3.3 Q(PDM) can be integrated into fabric by soaking with volatile organic solvents

EtOH solutions of q(PDM) were applied to general surgical facemasks and allowed to dry for at least 6 hrs at room temperature. 52% (S.D. 15%) of q(PDM) that was applied was integrated into the fabric of the mask. Polymer integration density (mg polymer / mm^2^ mask) was reported for each subsequent experiment and ranged between 0.01-0.12 mg/mm^2^.

PPE permeability after q(PDM) integration was also examined to ensure that the addition of polymer does not impact the breathability or permeability of the existing facemask. Upright cup tests revealed that exposure of EtOH inherently damaged the integrity of the facemask, however at densities above 0.02 mg/mm^2^ permeability was decreased below unmodified facemask controls (**Figure 6**). Subjectively, the decreased permeability of the facemask material did not significantly hinder breathability. We predict that administration of pure anhydrous EtOH dehydrated the fibers, causing the increased permeability seen in EtOH controls and the 0.01 mg/mm^2^ group. At higher integration densities, these ‘cracked’, dehydrated fibers are filled with q(PDM) upon solvent evaporation.

SEM images were obtained to ensure that the integrated q(PDM) was evenly distributed among the existing facemask fibers and showed that an increase in polymer integration yielded an increase in average fiber diameter size (**Figure 7**). At higher concentrations (>0.1 mg/mm^2^), droplets of solidified polymer were observed (**Figure S1**). Overall, q(PDM) integration with existing fibers was confirmed.

### 3.4 Facemasks with q(PDM) can prevent E. coli and S. aureus infiltration, even with imperfect fitting

‘Chronic exposure’ (10 minutes, nebulizer mist, ~5×10^8^ CFU/mL) trials showed that despite common belief, general surgical facemasks are permeable to bacteria over long-term exposure. **Figures 8** **and** **9** show that upon integration of 0.01 mg/mm^2^ q(PDM), bacterial filtration decreases, most likely due to the increased permeability caused by EtOH treatment. However, at higher concentrations (>0.05 mg/mm^2^) both live E. coli and S. aureus were either inactivated or filtered as shown on the (FT) dish, and colonies were unable to be cultured from the back (B) of each mask (**Figures 8** **and** **9**). As the integration density was increased to 0.12 mg/mm^2^, nebulized bacteria were significantly – or completely – eliminated from FT and B in both ‘acute’ and ‘chronic’ conditions. It was also observed that higher concentration q(PDM) facemasks were less likely to absorb condensate than traditional facemasks, most likely due to an increase in surface hydrophobicity.

### 3.5 Q(PDM) inhibits bacteria attachment and growth on facemask fabrics

Although bacterial survival on surfaces has been widely reported for both S. aureus (>7 days) and E.coli (>5 d) ^26–28^, we aimed to observe if both Gram-positive and Gram-negative bacteria can remain viable on a surgical facemask over the span of 16 hours after contamination, which we predicted would be the intraday time frame for exposure from handling a contaminated facemask. In both Gram-positive and Gram-negative trails, we observed that viable bacteria survived on the control and EtOH facemasks for up to 16 hours, however E. coli was unable to survive over 1 hour on facemasks with >0.04 mg/mm^2^ integrated polymer (**Figure 10a)** and S. aureus after 16 hours (**Figure 10b)**.

### 3.6 Q(PDM) can be electrospun into micro-scale fibers

Exploring alternative methods of integrating q(PDM) into existing fabrics and PPE, we investigated whether the polymer was able to be electrospun to produce homogenous, polymer ‘fiber mats’. Q(PDM) was dissolved in a variety of organic solvents (EtOH, DMF, THF) and at different weight percentages (5%, 10%, 15%). It was observed that EtOH was able to repeatedly dissolve q(PDM) at room temperature and was able to be fed through the electrospinning tubing without significant clotting (both DMF and THF were also compatible solvents, however more frequent clotting was observed). Electrospinning parameters, including solution flow rate and voltage potential, were varied, and it was found that ~10% w/w EtOH solution at 1mL/hr between 8-10 kV produced visible, white microfibers (**Table 1**). 5% w/w was also able to generate fibers, however light microscopy images revealed that the fibers were heterogeneously combined with polymer droplets, suggesting electrospraying was also occurring (**Figure S2**). 15% w/w caused clotting at the emitter and was unable to produce fibers. It should be noted that optimal parameters for electrospinning q(PDM) were dependent on batch and environmental conditions (e.g. temperature/humidity) and was adjusted accordingly until fiber formation was observed.

**Table 1:**
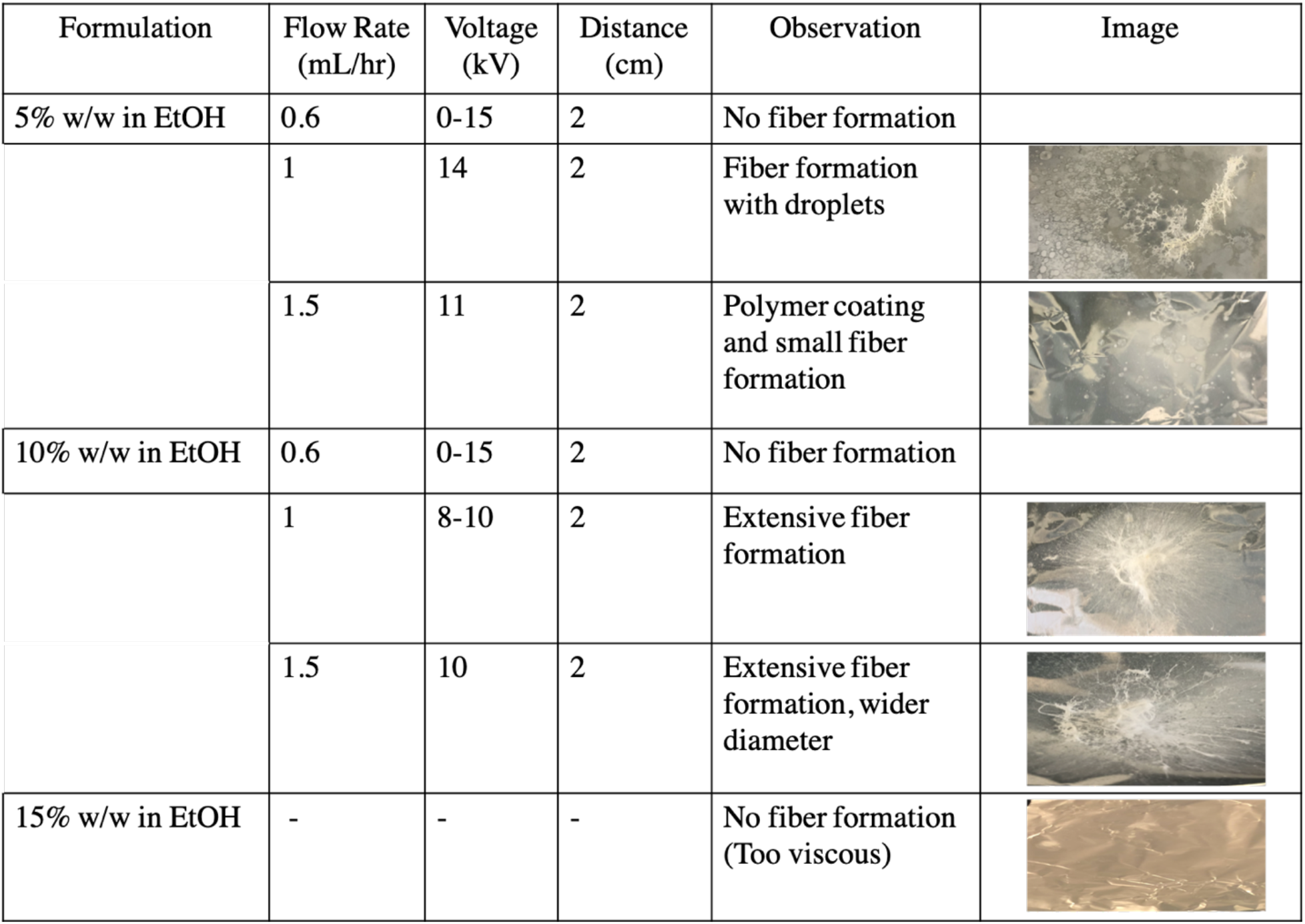
Electrospinning of q(PDM) with varied concentrations and electrospinning parameters.

| Formulation | Flow Rate (mL/hr) | Voltage (kV) | Distance (cm) | Observation | Image |
| --- | --- | --- | --- | --- | --- |
| 5% w/w in EtOH | 0.6 | 0-15 | 2 | No fiber formation |  |
|  | 1 | 14 | 2 | Fiber formation with droplets |  |
|  | 1.5 | 11 | 2 | Polymer coating and small fiber formation |  |
| 10% w/w in EtOH | 0.6 | 0-15 | 2 | No fiber formation |  |
|  | 1 | 8-10 | 2 | Extensive fiber formation |  |
|  | 1.5 | 10 | 2 | Extensive fiber formation, wider diameter |  |
| 15% w/w in EtOH | - | - | - | No fiber formation (Too viscous) |  |

Based on results in **Table 1**, narrower concentration ranges of q(PDM) were attempted, and it was wound that 7.5% w/w EtOH solutions produced fibers and prevented clotting in the electrospinner emitter. 7.5% w/w produced q(PDM) samples were then imaged via SEM and was observed to have fiber diameters ranging from 100nm - 100μm, with a notable ‘braided’ structure (**Figure 11**).

## 4. Discussions

Herein, we outlined a two-step synthesis of q(PDM), which was successfully polymerized into 9,800-30,000 g/mol polymer strands with ~75% quaternization (**Figure 4**). Changes in molecular weight and quaternization can be achieved by altering molar ratios, solvent amount, and time of reactions and, overall, our outlined method is relatively safe (i.e. common solvents, temperature not exceeding 70°C).

**Figure 4:**
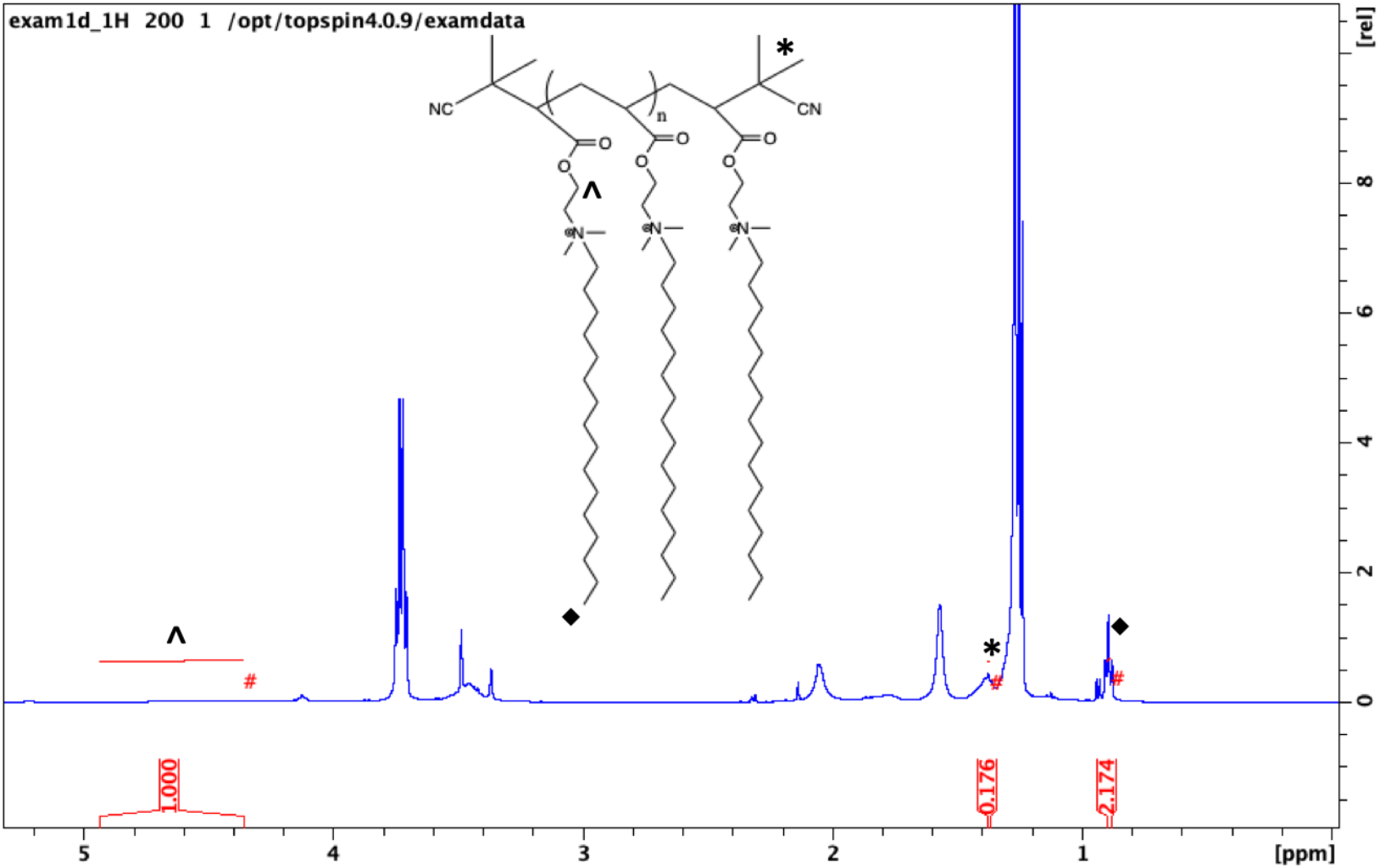
^1^H-NMR (DMSO-D_6_) of q(PDM), with unique peaks at δ =1.37 ppm (CH_3_ hexadecyl endgroup, ♦), δ = 0.88 ppm (AIBN terminal CH_3_, *), and δ = 4.52 ppm (CH_2_ neighboring DMAEMA’s ester group, ^).

The polymer was then integrated into surgical facemasks by soaking the existing fibers in EtOH solutions of q(PDM). Integration was confirmed by SEM, increasing existing fiber diameters by 7 μm (0.02 mg/mm^2^) up to over 40 μm (0.12 mg/mm^2^) (**Figure 7**, **S1**). As a proxy for breathability, mask permeability was studied to ensure that modified facemasks were still ‘breathable’. Mask permeability found to increase after treatment with EtOH, however sufficient addition of polymer (>0.02 mg/mm^2^) lowered permeability slightly below surgical facemask controls (**Figure 6**).

Antimicrobial properties of q(PDM) were then confirmed against both Gram-negative (E. coli) and Gram-positive (S. aureus) bacteria (**Figures 5**, **8**, **9**, **and** **10**). Broth-dilution assays revealed that q(PDM)’s antimicrobial properties are dependent on contact with bacteria and total polymer surface area (**Figure 5**). In a relatively high concentrations bacteria solution, it was observed that S. aureus (Gram-positive) was killed more efficiently than E. coli. (Gram-negative).

**Figure 5:**
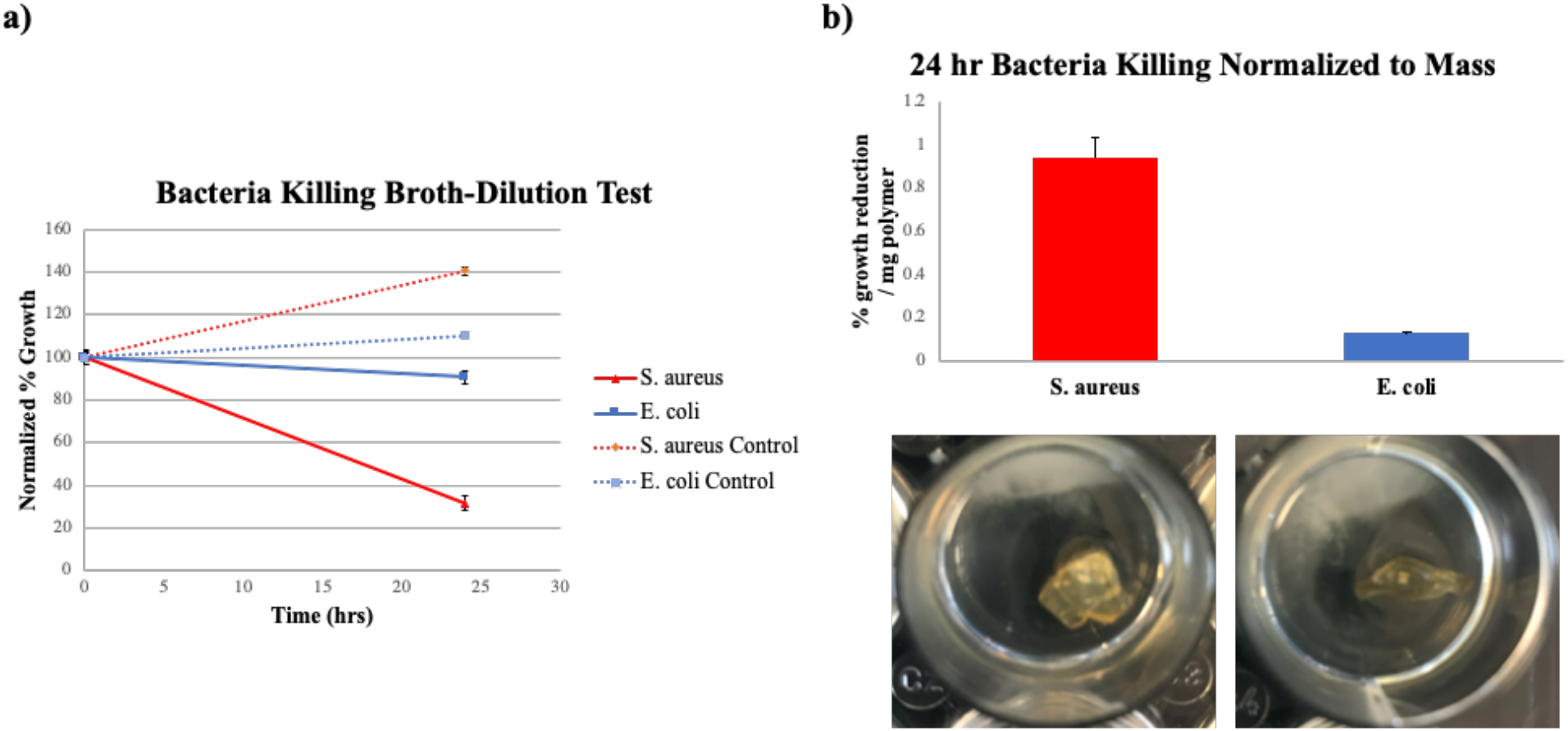
**a)** Growth reduction of bacteria after 24 hrs incubation with 40-50mg q(PDM) polymer, normalized to positive (bacteria-only) and negative (broth-only) controls. **b)** Percent growth inhibition normalized to polymer mass for both S. aureus and E. coli. In all groups (n=3), error bars are representative of the standard deviation.

**Figure 6:**
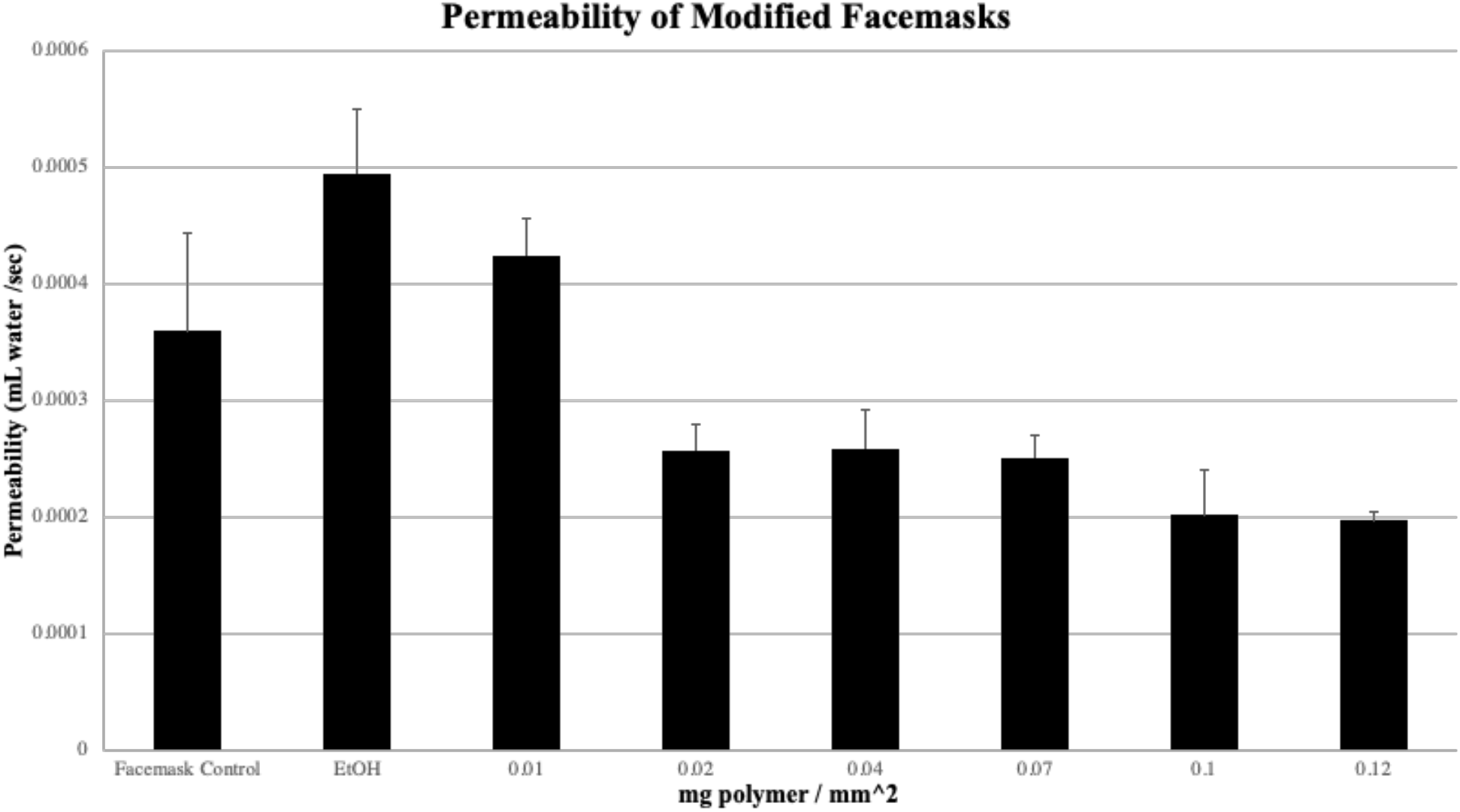
Upright cup test for facemasks integrated with q(PDM) EtOH solutions. Each group (n=3) was held at a constant 37°C, 18% humidity.

**Figure 7:**
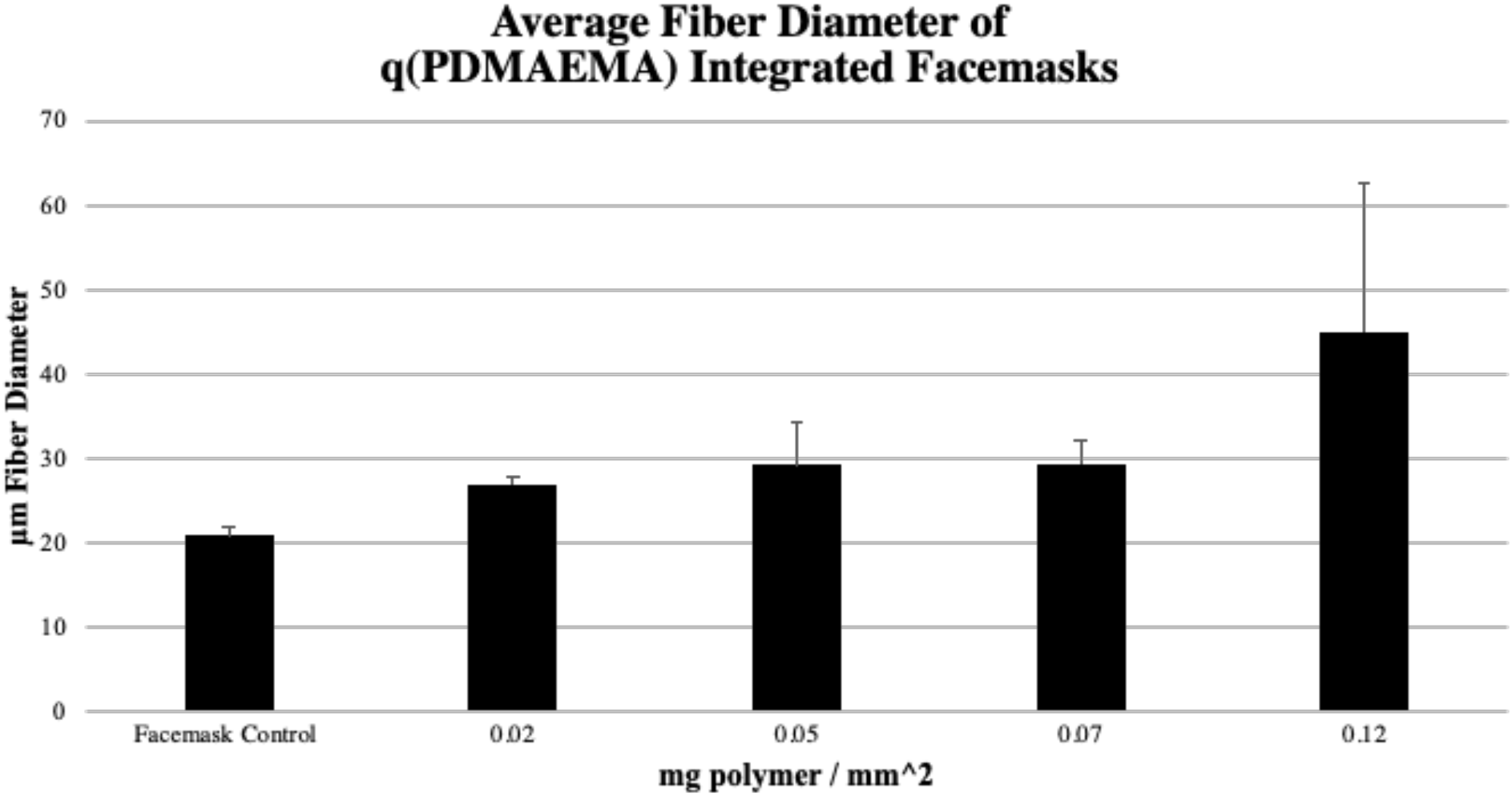
SEM image obtained average fiber diameter sizes of q(PDM) integrated facemasks (n=2). An average of 10 fiber diameters were measured per image (~20-30 total per group).

**Figure 8:**
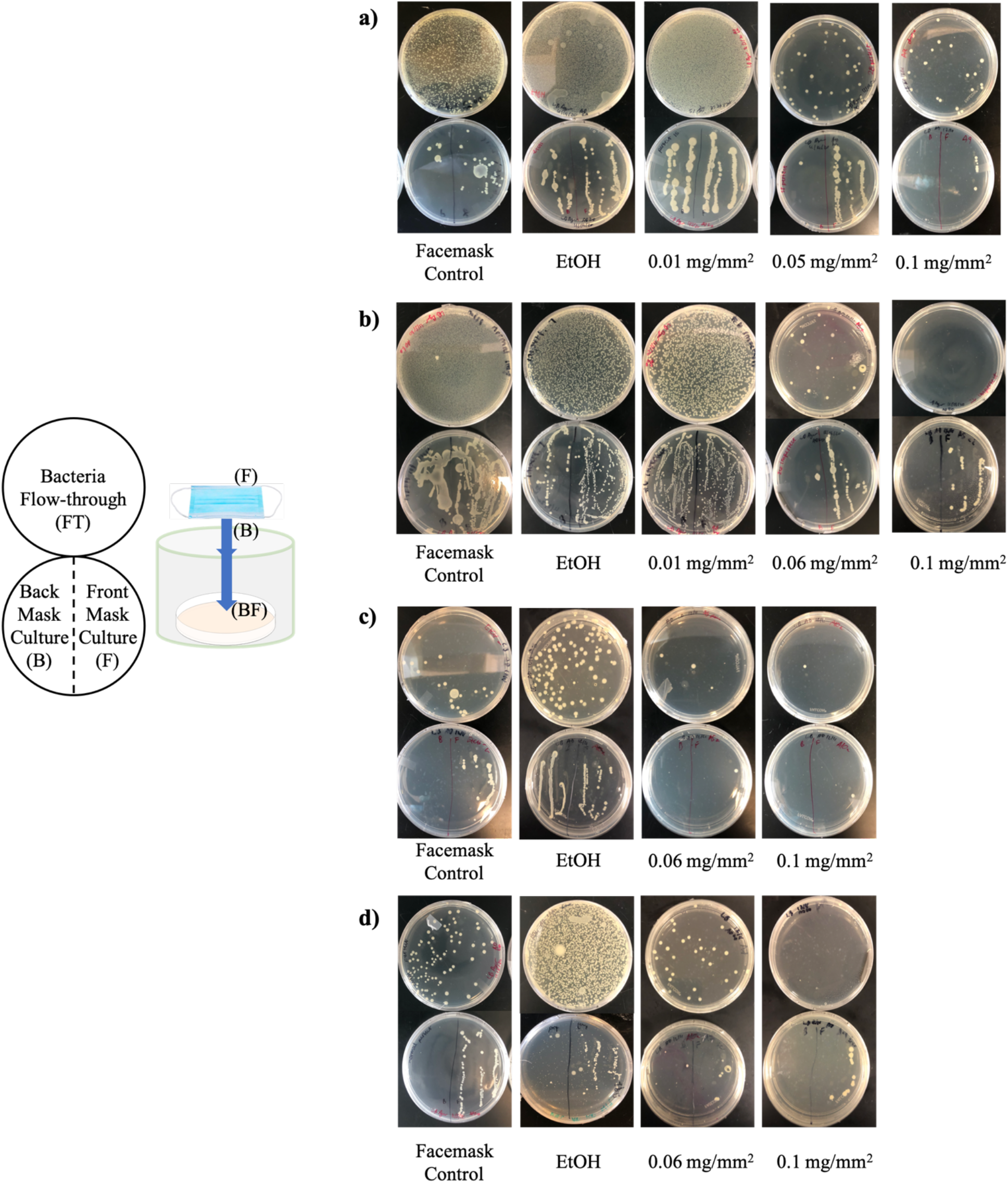
Nebulized E. coli ‘chronic exposure’ facemask test for **a)** ‘perfectly sealed’ facemasks and **b)** ‘imperfectly sealed’ facemasks (~5 holes). ‘Acute exposure’ (e.g. sneeze) **c)** ‘perfectly sealed’ facemasks and **b)** ‘imperfectly sealed’ facemasks.

**Figure 9:**
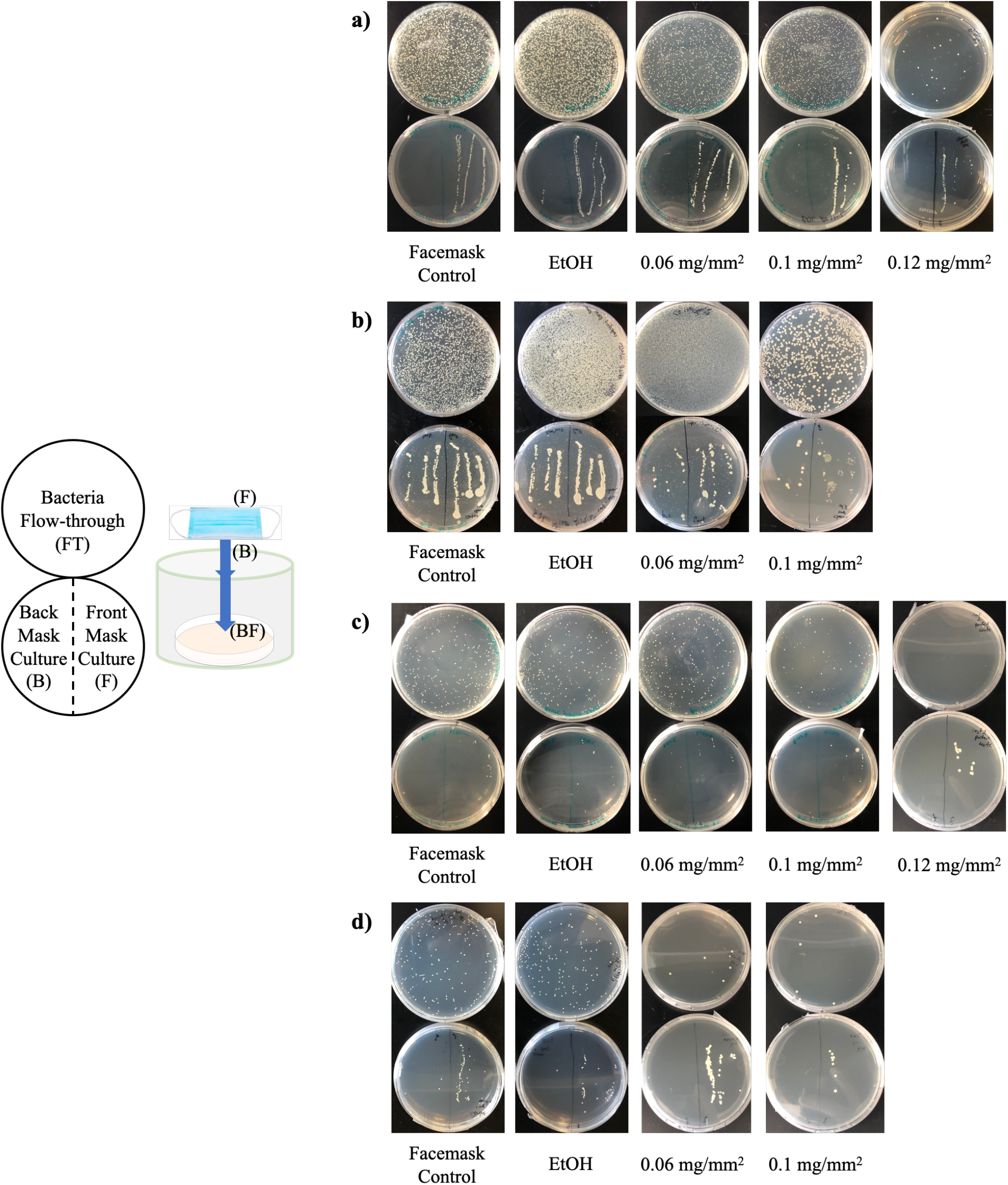
Nebulized S. aureus ‘chronic exposure’ facemask test for **a)** ‘perfectly sealed’ facemasks and **b)** ‘imperfectly sealed’ facemasks (~5 holes). ‘Acute exposure’ (e.g. sneeze) **c)** ‘perfectly sealed’ facemasks and **b)** ‘imperfectly sealed’ facemasks

**Figure 10:**
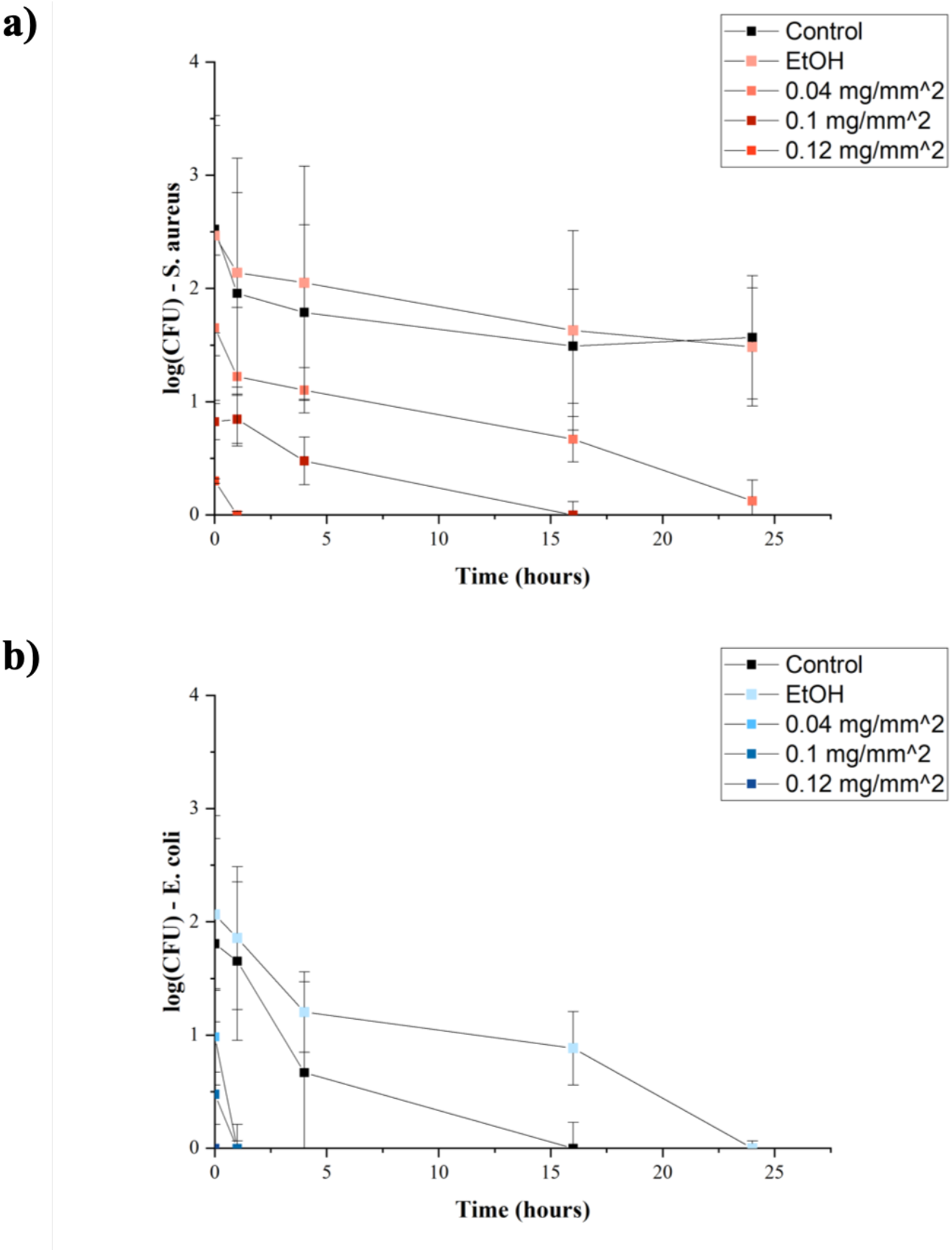
Surface contact-killing assay for both **a)** S. aureus and **b)** E. coli. Error bars are representative of standard deviation of n=3 trials.

Once the polymer was confirmed to have antimicrobial properties, aerosolized bacteria and low-concentration bacteria contact-killing assays were used to better simulate facemask contamination conditions *in vitro* (i.e. droplet exposure, ‘sneezes’). Both Gram-negative and Gram-positive aerosolized bacteria were inactivated/filtered in “chronic” and “acute” conditions with q(PDM)-integrated facemasks (**Figures 8** **and** **9**). Even in simulated “imperfect fits” where masks were damaged prior to bacteria exposure, the total, viable bacterial load was greatly deceased (**Figures 8b,d** **and** **9b,d**). Contrary to bulk broth dilution tests, it was found that nebulized E. coli was more efficiently filtered/inactivated compared to S. aureus. Similarly, bacteria contact-killing assays revealed that the survival time of E. coli was reduced to only 1 hour on q(PDM) facemasks, while S. aureus was able to persist until around 16 hours (**Figure 10**).

Differences in bactericidal activity are most likely attributed to differences in E. coli and S. aureus cellular structures: Gram-positive S. aureus have significantly thicker peptidoglycan cell walls (~50 nm) compared to Gram-negative E. coli’s inner/outer lipid membranes (~2 nm)^29^. Since quaternary ammonium polymers act by disrupting bacterial membranes, we believe that while E. coli can survive better as a biofilm, as observed in the broth-dilution assay, they are more easily killed in aerosolized-form or low-concentration solutions. Conversely, S. aureus is able to survive more effectively in aerosolized form and at low-concentrations as its peptidoglycan cell walls are more resistant towards q(PDM)’s active moieties, and therefore requires higher polymer concentrations to effectively inactivate.

We then explored alternate processing of q(PDM) via electrospinning. Electrospinning, and newer methods including melt electrospinning^30^, are scalable manufacturing processes used to efficiently produce micro- and nanoscale fiber networks from polymers^31^. By adjusting concentration, voltage, flow rate, and collecting distance, we determined that a 7.5% w/w solution of q(PDM) in EtOH produced electrospun fibers with diameters between 100 nm to 100 μm (**Table 1**, **Figure 11**).

**Figure 11:**
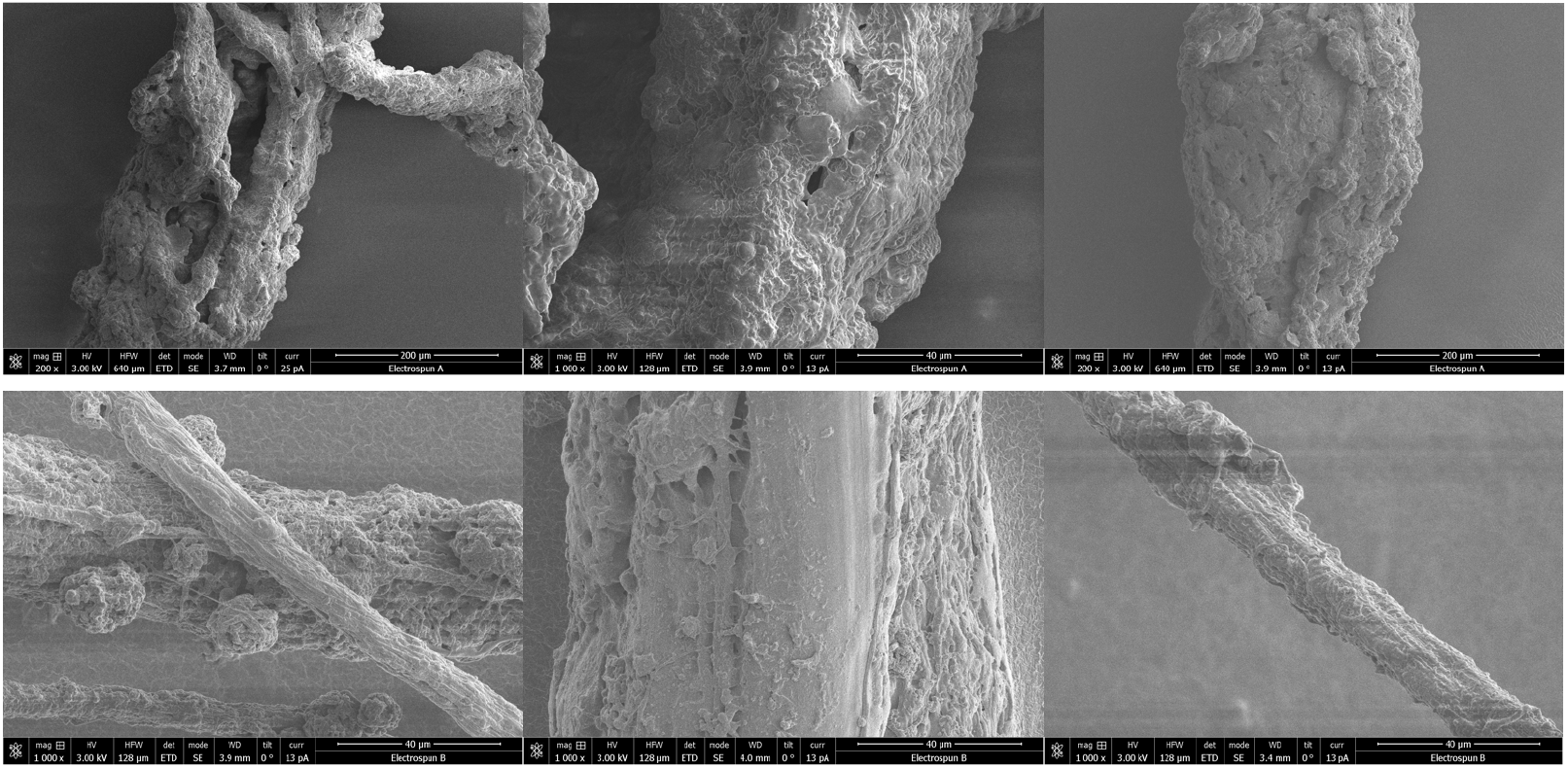
SEM images of q(PDM) electrospun fibers from 7.5% w/w EtOH solution. Fiber diameter sizes ranged from 100nm - 100μm.

While not included in this study, future studies will investigate the antimicrobial activity of q(PDM) electrospun mats. High volume electrospinning, for fiber mat production and manufacturing, will require additional parameter optimization; however, electrospun mats have generally been shown to effectively filter aerosolized particles ^32–34^. In addition, based on the proposed cytoplasmic membrane disrupting mechanism of alkane-branched quaternary cationic polymers, q(PDM) may have viricidal against enveloped viruses, which will also be a focus of future studies^35,36^.

## 5. Conclusion

To our knowledge, this is the first integration of q(PDM) for the purpose of generating antimicrobial fabrics and personal protective equipment. Herein, we outlined a method to synthesize a quaternary ammonium polymer, q(PDM), coated existing PPE fabric fibers with q(PDM) and generated microfibers of pure q(PDM) via electrospinning. q(PDM) was observed to be effective against both Gram-positive and Gram-negative bacteria and can significantly decrease survival time of bacteria (both in broth and aerosolized) on its surface. This study presents q(PDM) as a potential solution for generating antiseptic personal protective equipment at a relatively low cost (manufactured at <$0.50 per gram polymer). Furthermore, q(PDM)’s hypothesized virucidal activity makes it a promising additive to PPE to counter airborne pathogens, including viruses.

## Author Contributions

H.A.R. and A.B.D. devised the project. A.B.D. developed the technical procedures and performed the experiments under the supervision of H.A.R. K.D. helped perform nebulizer assays. Under the supervision of H.A.R, A.B.D wrote the manuscript; all authors read or edited the manuscript.

## Acknowledgements

Special thanks to the Advanced Platform Technology Center of the Louis Stokes VA Cleveland Medical Center for funding this project and the Swagelok Center for Surface Analysis of Materials for the use of their Helios SEM.

---

**S1.**
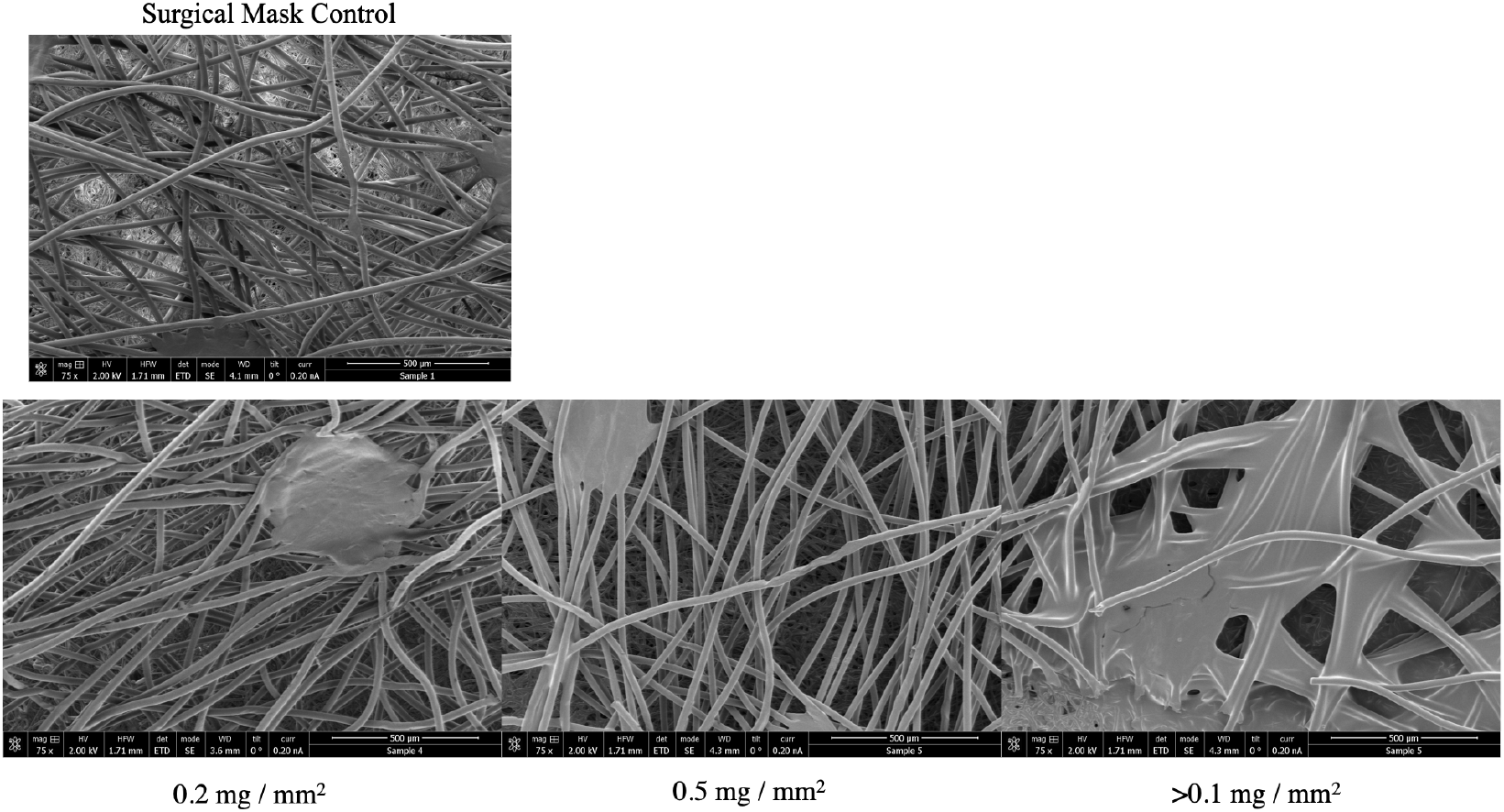
SEM images of facemasks integrated with q(PDM) at various concentrations

**S2.**
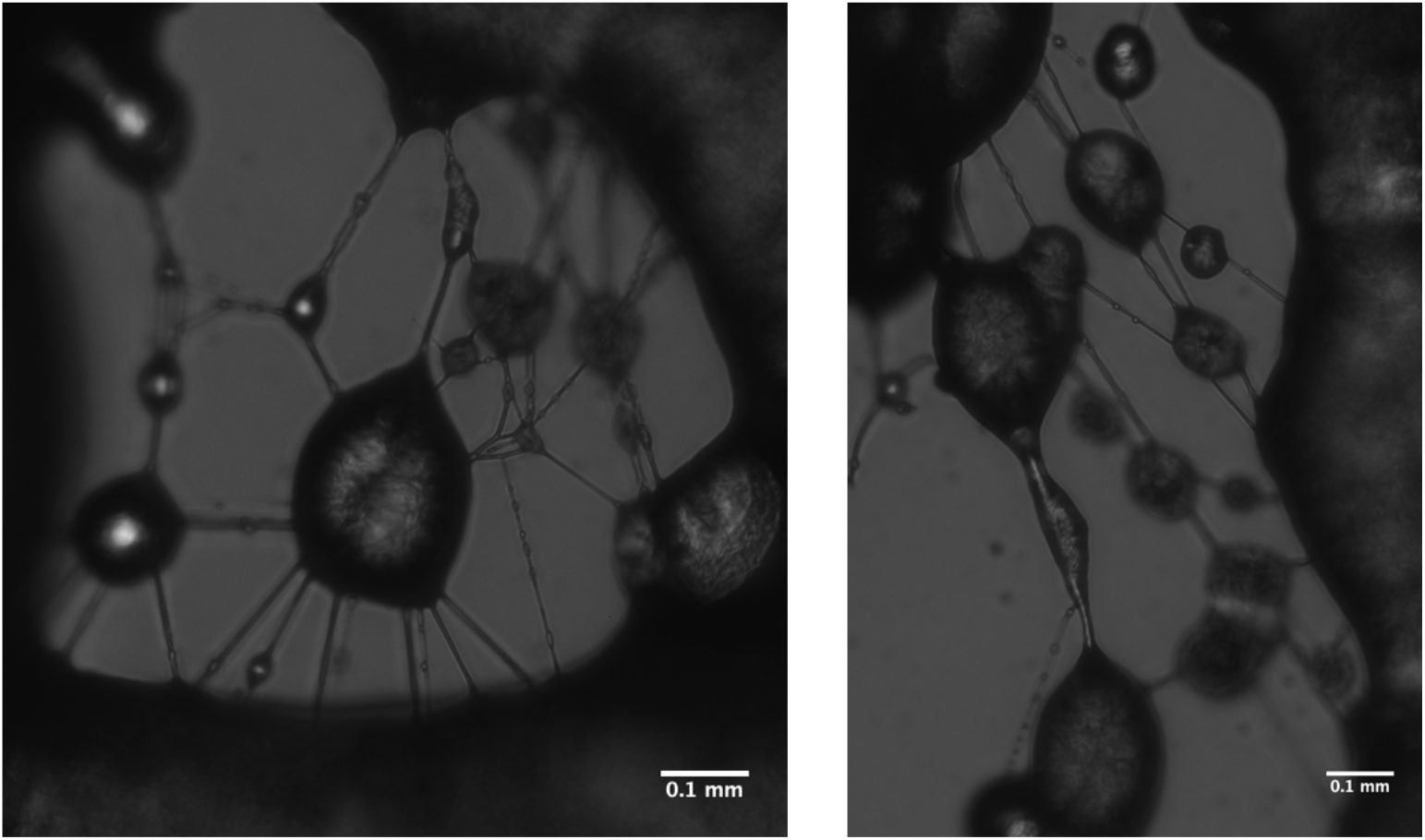
Light microscopy images of 5% w/w EtOH electrospun fibers at 1 mL/hr, 10 kV.

