## Supplementary material for "Polymer additives to personal protective equipment can inactivate pathogens": Figure S1

Surgical Mask Control

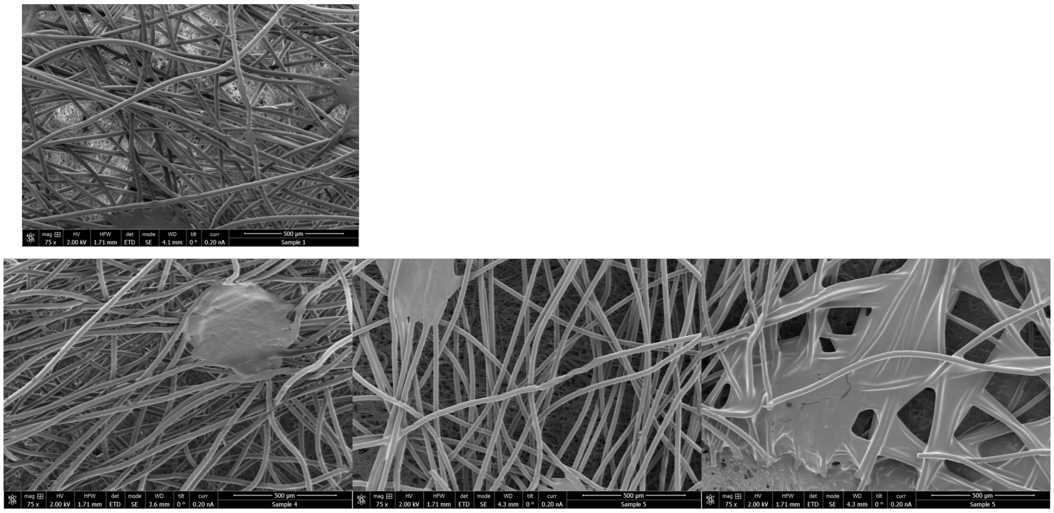

0.2 mg / mm²     0.5 mg / mm²     >0.1 mg / mm²

**S1**. SEM images of facemasks integrated with q(PDMAEMA) at various concentrations
