## Supplementary figures and images for "Polymer additives to personal protective equipment can inactivate pathogens"

### Figure S2

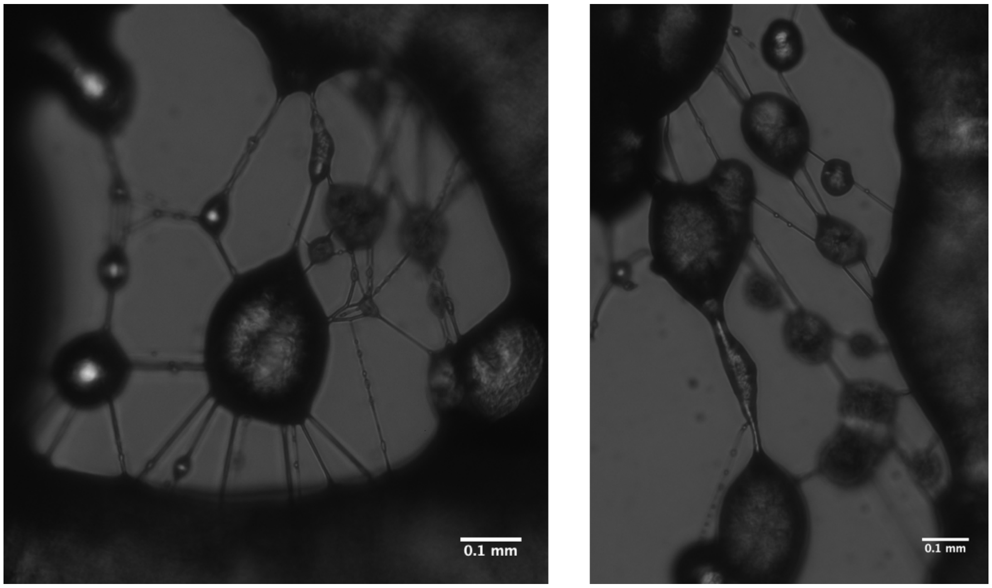


**S2**. Light microscopy images of 5% w/w EtOH electrospun fibers at 1 mL/hr, 10 kV.
